# Back to sequences: find the origin of *k*-mers

**DOI:** 10.1101/2023.10.26.564040

**Authors:** Anthony Baire, Pierre Peterlongo

## Abstract

A vast majority of bioinformatics tools dedicated to the treatment of raw sequencing data heavily use the concept of *k*-mers. This enables us to reduce the data redundancy (and thus the memory pressure), to discard sequencing errors, and to dispose of objects of fixed size that can be manipulated and easily compared to each others. A drawback is that the link between each *k*-mer and the original set of sequences it belongs to is generally lost. Given the volume of data considered in this context, finding back this association is costly. In this work, we present “`back_to_sequences`”, a simple tool designed to index a set of *k*-mers of interests, and to stream a set of sequences, extracting those containing at least one of the indexed *k*-mer. In addition, the number of occurrences of *k*-mers in the sequences is provided. Our results show that `back_to_sequences` streams *≈* 200 short read per millisecond, enabling to search *k*-mers in hundreds of millions of reads in a matter of a few minutes.

**Availability:** github.com/pierrepeterlongo/back_to_sequences

## 1 Introduction

In the 2010s, following the emergence of next-generation sequencing technology, the read assembly strategies based on the overlap-layout-consensus (OLC) paradigm were unable to scale to tens of millions of reads or more, prompting the usage of the *de Bruijn* graph (dBG) data structure [4, 14]. The dBG success was due to the fact that the main difficulties associated with the sequencing data nature (read redundancy, non-uniform coverage and non-uniform overlap between reads, sequencing errors, unknown sequencing strand) were complex to handle with the OLC while being easy to handle or simply solved with the dBG approach [10].

Recall that in the dBG assembly approach, 1. all *k*-mers (words of length *k*) from a set of reads are counted; 2. those with an abundance lower than a threshold are considered as containing sequencing errors and are discarded; 3. the remaining *k*-mers are organized in a dBG; 4. Paths from the dBG form the basis of the assembly, later improved thanks to scaffolding tools [7] such as tools provided for instance by the Spades assembler [1].

The *k*-mers usefulness did not end with their use in dBGs. A large and redundant set of sequences such as a sequencing read set, can be summarized to its set of *k*-mers. Among multiple fundamental tasks, it has been the basis for metagenome comparisons [2], for taxonomy characterization [17], for indexing purposes [3, 9], for genotyping [6], for identification [13], for transcript expression estimation [18], or for variant discovery [16] to cite only a few.

One of the keys to the success of the usage of *k*-mers stands in their low resource needs. Indeed whatever the sequencing coverage, once filtered, the number of distinct *k*-mers is roughly at most equal to the original genome size. This offers a low impact on RAM and/or disk needs. However, this comes at the cost of losing the link between each kmer and the sequences(s) from which it originates. Storing explicitly these links would reintroduce the problem associated with the abundance of original reads, as the link between each original read and each of its *k*-mers would have to be stored.

Finding back the link between a *k*-mer and each original read in which it occurs can be performed by indexing the reads [11] which hardly scales hundreds of million reads. One may also apply *grep-like* evolved pattern matching approaches as `pt` [12]. However, even if they were highly optimized these last decades, these approaches cannot detect efficiently thousands of *k*-mers in millions of reads. The approaches that use *k*-mers for genotyping such as `kage` [6] may find the number of occurrences of the *k*-mers but not extract the sequences from which they originate.

In this context, we propose a simple tool, named `back_to_sequences`, specifically dedicated to extracting from a set of sequences (eg. reads), those that contain some of the *k*-mers given as input and counting the number of occurrences of each of such *k*-mers in the sequences. The *k*-mers can be considered in their original, reverse-complemented, or canonical form.

## 2 Results

The tool we introduce, `back_to_sequences`, uses the native `rust` data structures to index and query *k*-mers. Its advantages are 1. its simplicity (install and run); 2. its low RAM usage; 3. its fast running time; 4. its additional features adapted to *k*-mers: control of their orientation (canonical, original or reversed-complemented) and sequence filtration based on the minimal and maximal percent of *k*-mers shared with the indexed set. To the best of our knowledge, there exists no other tool dedicated to this specific task.

We propose a simple performance benchmark to assess the `back_to_sequences` scaling performances on random sequences and on real metagenomic data.

### 2.1 Benchmark on random sequences

We generated a random sequence *S* on the alphabet {*A, C, G, T*} of size 100 million base pairs. From *S*, we randomly extracted 50,000 sub-sequences each of size 50. We consider these sequences as the set of *k*-mers to be searched. As we used *k* = 31, each subsequence contains 50 − 31 + 1 = 20*kmers*. Doing so, we consider a set of at most 50000 × 20 = 1, 000, 000 *k*-mers. We call those *k*-mers the “reference-*k*-mers”. Some of them can be redundant.

We also generated six sets: {*Q*_10*k*_, *Q*_100*k*_, *Q*_1*M*_, *Q*_10*M*_, *Q*_100*M*_, *Q*_200*M*_}, composed respectively of 10 thousand, 100 thousand, one million, 10 million, 100 million, and 200 million sequences, each of length 100 nucleotides. Each sequence is randomly sampled from *S*.

In each sequence of each set *Q*_*i*_, we searched for the existence of *k*-mers indexed in the “reference-*k*-mers” set. The performances are provided Table 1. Presented results were obtained on the GenOuest platform on a node with 32 threads Xeon 2.2 GHz, and on a MacBook, Apple M2 pro, 16 GB RAM with 10 threads, respectively denoted by “Time GenOuest” and “Time mac” in this table. These results highlight the scalability of the `back_to_sequences` tool, able to search a million of *k*-mers in hundreds of millions of reads on a laptop computer in a matter of dozens of minutes with negligible RAM usage.

**Table 1:** The back_to_sequences performances searching one millions *k*-mers in 10 thousands to 100 million reads. Tested version: 2.5.0.

| Number of reads | Time GenOuest | Time mac | max RAM |
| --- | --- | --- | --- |
| 10,000 | 0.9s | 0.6s | 0.13 GB |
| 100,000 | 1.0s | 1.4s | 0.13 GB |
| 1,000,000 | 5.3s | 8.3s | 0.13 GB |
| 10,000,000 | 31s | 39s | 0.13 GB |
| 100,000,000 | 5m39 | 6m04 | 0.13 GB |
| 200,000,000 | 10m40 | 12m57 | 0.13 GB |

### 2.2 Benchmark on *Tara* ocean seawater metagenomic data

The previously proposed benchmark shows the scalability of the proposed approach. Although performed on random sequences, there is no objective reason why performance should differ on real data, regardless of the number of *k*-mers actually detected in the data. We verify this claim by applying `back_to_sequences` to real complex metagenomic sequencing data from the *Tara* ocean project [15].

We downloaded one of the *Tara* ocean read sets: station number 11 corresponding to a surface Mediterranean sample, downloaded from the European Nucleotide Archive, identifier ERS488262^1^. We extracted the first 100 million reads, which are all of length 100. Doing so, this test is comparable to the result presented Table 1 querying 100 million random reads. Using `back_to_sequences` we searched in these reads each of the 69 31-mers contained in its first read. On the GenOuest node, `back_to_sequences` enabled to retrieve all reads that contain at least one of the indexed *k*-mers in 5m17 with negligible RAM usage of 45MB. As expected, these scaling results are in line with results presented Table 1.

Out of curiosity, we ran `back_to_sequences` on the full read set, composed of ≈ 26.3 billion *k*-mers, and 381 million reads, again for searching the 69 *k*-mers contained in its first read. This operation took 20m11.

### 2.3 Possible alternatives

To the best of our knowledge, there exists no tool specifically dedicated to finding sequences that contain at least one *k*-mer among a set.

Genotypers as `kage` [6], using `kmer mapper` [8], provide a way to count the number of occurrences of each *k*-mer from a set of reference *k*-mers in a read file. However, they do not offer the feature to extract reads that contain any reference *k*-mer.

Finding one unique *k*-mer of interest in a set of sequences can be done using the classical `grep` or more recent pattern matching tools such as “*The Platinum Searcher*” [12] or “*The Silver Searcher*” [5].

As for testing, on the MacBook, Apple M2 pro, we queried one *k*-mer in the *Q*_100*M*_ dataset (see Section 2.1) using `grep`, *The Platinum Searcher*, and *The Silver Searcher*.

- `grep` required 44 seconds. Thus, by simple extrapolation, searching for one million *k*-mers on a single computer would require approximately 500 days, to be compared to 5 to 6 minutes using `back_to_sequences`.
- `pt` (*The Platinum Searcher*) required 15 seconds, which can be extrapolated to approximately 175 days if searching for one million *k*-mers.
- `ag` (*The Silver Searcher*) did not finish after 400 seconds.

Summing up, these alternative tools are not meant for querying numerous patterns at the same time and do not scale to large problem instances.

Note also that these alternative tools are not specialized to genomic data in which one is interested in searching for a *k*-mer and potentially its reverse complement. Finally, these tools do not easily provide the number of occurrences of each of the searched patterns when they are many.

## 3 Method and features

As previously stated, `back_to_sequences` is a very simple tool, written in `rust`. It uses the native HashMap for storing the searched *k*-mer set. Depending on the user choice, the original or the canonical (the smallest alphabetical word between the *k*-mer and its reverse complement) version of each *k*-mer from the “reference-*k*-mers” set is indexed.

At query time, given a sequence *s*, all its *k*-mers are extracted and queried. Depending on the user’s choice, the canonical or the original representation of each *k*-mer from the reference set is indexed. Again, depending on the user’s choice, for each queried *k*-mer, the original version, its reverse complement, or its canonical representation is queried. If the queried *k*-mer belongs to the index, its number of occurrences is increased. The number of *k*-mers extracted from the sequence *s* that have a match with the index is output together with the original sequence. Sequences are queried in parallel.

A minimal and a maximal threshold can be fixed by the user. A queried sequence is output only if its percentage of *k*-mers that belong to the searched *k*-mers is strictly higher than the minimal threshold and lower or equal to the maximal threshold. While the minimal threshold enables to focus on sequences that are similar enough with a set of *k*-mers, the maximal threshold offers for instance a way to remove contaminated sequences.

The sequences to be queried can be provided as a fast or fastq file (gzipped or not). They can also be be read directly from the standard input (*stdin*). This offers the may to stream sequences as they arrive, for instance when they are obtained during an Oxford Nanopore sequencing process.

The output format of queried sequences respects the input format: if the input is in fasta (resp. fastq), the output is in fasta (resp. fastq).

## 4 Conclusion

We believe that `back_to_sequences` is a generic and handy tool that will be beneficial for building pipelines that require manipulating *k*-mers and finding back the sequences from which they originate and/or counting their number of occurrences in a set of genomic sequences. We also believe that `back_to_sequences` will have other straightforward applications such as quality control, contamination removal, or genotyping known pieces of sequences in raw sequencing datasets, all of which being possible in real-time throughout the sequencing process.

## Acknowledgements

We acknowledge the GenOuest core facility (https://www.genouest.org) for providing the computing infrastructure.

## Footnotes

1 http://www.ebi.ac.uk/ena/data/view/ERS488262

## Notes

https://github.com/pierrepeterlongo/back_to_sequences

### Competing Interest Statement

The authors have declared no competing interest.

## References

[1] Anton Bankevich, Sergey Nurk, Dmitry Antipov, Alexey A Gurevich, Mikhail Dvorkin, Alexander S Kulikov, Valery M Lesin, Sergey I Nikolenko, Son Pham, Andrey D Prjibelski, et al. Spades: a new genome assembly algorithm and its applications to single-cell sequencing. Journal of computational biology, 19(5):455–477, 2012.

[2] Gaëtan Benoit, Pierre Peterlongo, Mahendra Mariadassou, Erwan Drezen, Sophie Schbath, Dominique Lavenier, and Claire Lemaitre. Multiple comparative metagenomics using multiset k-mer counting. PeerJ Computer Science, 2:e94, 2016.

[3] Andrea Cracco and Alexandru I Tomescu. Extremely fast construction and querying of compacted and colored de bruijn graphs with ggcat. Genome Research, pages gr–277615, 2023.

[4] Paul Flicek and Ewan Birney. Sense from sequence reads: methods for alignment and assembly. Nature methods, 6(Suppl 11):S6–S12, 2009.

[5] Geoff Greer. The Silver Searcher. https://github.com/ggreer/the_ silver_searcher, 2020. [Online; accessed 24-October-2023].

[6] Ivar Grytten, Knut Dagestad Rand, and Geir Kjetil Sandve. Kage: Fast alignment-free graph-based genotyping of snps and short indels. Genome Biology, 23(1):209, 2022.

[7] Daniel H Huson, Knut Reinert, and Eugene W Myers. The greedy path-merging algorithm for contig scaffolding. Journal of the ACM (JACM), 49(5):603–615, 2002.

[8] Knut Dagestad Rand Ivar Grytten. Kmer Mapper. https://github.com/ivargr/kmer_mapper, 2020. [Online; accessed 24-October-2023].

[9] Téo Lemane, Paul Medvedev, Rayan Chikhi, and Pierre Peterlongo. Kmtricks: efficient and flexible construction of bloom filters for large sequencing data collections. Bioinformatics Advances, 2(1):vbac029, 2022.

[10] Zhenyu Li, Yanxiang Chen, Desheng Mu, Jianying Yuan, Yujian Shi, Hao Zhang, Jun Gan, Nan Li, Xuesong Hu, Binghang Liu, et al. Comparison of the two major classes of assembly algorithms: overlap–layout–consensus and de-bruijn-graph. Briefings in functional genomics, 11(1):25–37, 2012.

[11] Camille Marchet, Lolita Lecompte, Antoine Limasset, Lucie Bittner, and Pierre Peterlongo. A resource-frugal probabilistic dictionary and applications in bioinformatics. Discrete Applied Mathematics, 274:92–102, 2020.

[12] Monochromegane. The Platinum Searcher. https://github.com/monochromegane/the_platinum_searcher, 2018. [Online; accessed 24-October-2023].

[13] Shahab Sarmashghi, Kristine Bohmann, M Thomas P. Gilbert, Vineet Bafna, and Siavash Mirarab. Skmer: assembly-free and alignment-free sample identification using genome skims. Genome biology, 20:1–20, 2019.

[14] Michael C Schatz, Arthur L Delcher, and Steven L Salzberg. Assembly of large genomes using second-generation sequencing. Genome research, 20(9):1165–1173, 2010.

[15] Shinichi Sunagawa, Silvia G Acinas, Peer Bork, Chris Bowler, Tara Oceans Coordinators, Damien Eveillard, Gabriel Gorsky, Lionel Guidi, Daniele Iudicone, Eric Karsenti, Fabien Lombard, Hiroyuki Ogata, Stephane Pesant, Matthew B Sullivan, Patrick Wincker, and Colomban de Vargas. Tara oceans: towards global ocean ecosystems biology. Nat Rev Microbiol, 18(8):428–445, 2020.

[16] Raluca Uricaru, Guillaume Rizk, Vincent Lacroix, Elsa Quillery, Olivier Plantard, Rayan Chikhi, Claire Lemaitre, and Pierre Peterlongo. Reference-free detection of isolated snps. Nucleic acids research, 43(2):e11–e11, 2015.

[17] Derrick E Wood, Jennifer Lu, and Ben Langmead. Improved metagenomic analysis with kraken 2. Genome biology, 20:1–13, 2019.

[18] Zhaojun Zhang and Wei Wang. Rna-skim: a rapid method for rna-seq quantification at transcript level. Bioinformatics, 30(12):i283–i292, 2014.

